# Enteropathy-induced regulatory T cells inhibit intestinal CD4+ T cell responses against oral vaccines

**DOI:** 10.1101/2020.06.03.130831

**Authors:** Amrita Bhattacharjee, Ansen H.P. Burr, Abigail E. Overacre-Delgoffe, Justin T. Tometich, Deyi Yang, Brydie R. Huckestein, Jonathan L. Linehan, Sean P. Spencer, Jason A. Hall, Oliver J. Harrison, Denise Morais da Fonseca, Elizabeth B. Norton, Yasmine Belkaid, Timothy W. Hand

**Affiliations:** R.K. Mellon Institute for Pediatric Research, Pediatrics Department, Infectious Disease Section, UPMC Children’s Hospital of Pittsburgh, University of Pittsburgh, Pittsburgh PA, 15224; Program in Microbiology and Immunology, University of Pittsburgh; Central South University, Xiangya School of Medicine, Changsha, Hunan, China; Metaorganism Immunity Section, Laboratory of Immune System Biology, National Institute of Allergy and Infectious Disease, National Institutes of Health, Bethesda MD; Department of Microbiology and Immunology, Tulane University School of Medicine, New Orleans, LA; Genentech Inc., South San Francisco, CA; Department of Microbiology and Immunology, Stanford University School of Medicine, Palo Alto, CA; ArsenalBio Inc. San Francisco, CA; Benaroya Research Institute, Seattle WA; Department of Immunology, Institute of Biomedical Sciences, University of São Paulo, Brazil

## Abstract

Environmental Enteric Dysfunction (EED) is an intestinal disease caused by malnutrition and infection that leads to malabsorption and stunting. EED is also associated with a reduced efficacy of oral vaccines. We show in a microbiota and diet-dependent model of EED that oral vaccine-specific CD4^+^ T cell responses fail in the small intestine but responses in the draining lymph node were unaffected. Accordingly, the number of immunosuppressive RORγT^+^FOXP3^+^ T_regs_ in the small intestine inversely correlated with the response to oral vaccination. Depletion of RORγT^+^FOXP3^+^ T_regs_ indicated that they were necessary for EED-associated inhibition of the vaccine response. Additionally, RORγT^+^FOXP3^+^ T_regs_ are important to regulate EED-associated inflammation as their depletion significantly worsened stunting. We have shown that EED-associated intestinal inflammation leads to a localized intestinal blockade of CD4 T cell immunity. These results support a modular model for immunity where tissue responses can be regulated independently of systemic immunity to prevent autoinflammation.

## Introduction

In the intestine, the immune system must balance the necessity for protection against infectious microorganisms while avoiding constant activation by the resident microbiome, which can lead to autoinflammatory disorders such as Inflammatory Bowel Disease. This task is made more difficult by the ever-changing environment of the intestine, where the host/microbiome relationship can be modified by shifts in diet and infection (Carmody et al., 2015; Cotillard et al., 2013; Hand, 2016). Diet-driven shifts to the microbiome have been associated with a variety of chronic diseases, and it is now believed that therapies that restore health to the microbiome will be critical for effective treatments (Blanton et al., 2016; Hand et al., 2016; Ridaura et al., 2013; Turnbaugh et al., 2006). Regardless of the cause, dysbiosis of the microbiome affects the immune response both locally, at barrier sites, and systemically, contributing to inflammatory disease (Henao-Mejia et al., 2012; Honda and Littman, 2016).

Oral vaccines are an effective mechanism to provide immunity to infection in the gastrointestinal tract and have been used to prevent the spread of enteric viral and bacterial infections (Vela Ramirez et al., 2017). Bacterial enterotoxins, such as the *E. coli* heat labile toxin (LT), have the potential to be excellent oral vaccines because they have intrinsic adjuvant activity and attenuated enterotoxins are currently in trials as vaccines against bacterial diarrhea or adjuvants for other pathogens (Clements and Norton, 2018; Harro et al., 2019; Norton et al., 2011; Shaikh et al., 2020). Oral vaccines can induce long-lived intestinal-resident CD4^+^ T cells and IgA^+^ plasma B cells, which mediate protection at the site of infection (Bemark et al., 2016; Clements and Norton, 2018; Pasetti et al., 2011). How intestinal-resident T and B cells develop and are regulated after oral vaccination is not completely understood but is important to our ability to design more effective vaccines that provide long-lived protection.

Regulatory T cells (T_regs_) are important for the maintenance of systemic and mucosal immune homeostasis(Josefowicz et al., 2012; Kim et al., 2016; Singh et al., 2001). One can broadly categorize intestinal T_regs_ responding to antigens derived from either self, food or the microbiota based on Neuropilin (NRP1) and RAR–related orphan receptor γ 2(RORγT) expression, where T_regs_ responding to self-antigens are NRP1^+^ RORγT^-^, those responding to microbiome-derived antigens are NRP1^-^ RORγT^+^ and those responding to food are negative for both molecules (Kim et al., 2016; Ohnmacht et al., 2015; Sefik et al., 2015; Weiss et al., 2012; Xu et al., 2018). In the intestine, both RORγT^+^ T helper 17 (T_h17_) cells and RORγT^+^FOXP3^+^ T_regs_ respond to antigens derived from intestinal adherent bacteria such as *Akkermansia spp*. and *Helicobacter spp*. or from bacteria that secrete antigen-bearing endosomes (Ansaldo et al., 2019; Chai et al., 2017; Kim et al., 2018; Sefik et al., 2015; Wegorzewska et al., 2019; Xu et al., 2018). The signals that determine whether T_reg_ or T helper cells dominate in the response to a given intestinal-resident bacterium are not completely known and may be contextual to environmental factors beyond the biology of antigen-bearing organisms (Ansaldo et al., 2019; Chiaranunt et al., 2018; Hand et al., 2012; Zhao et al., 2017). The balance of T_regs_ and T_h_ cells in the intestine is important because inflammatory diseases have been associated with an increased frequency of T_h1_/T_h17_ cells and decreased T_regs_ (Leung et al., 2014; Lochner et al., 2008). T_regs_ also support the generation of IgA-producing plasma cells, particularly within Peyer’s Patches (Cong et al., 2009; Kawamoto et al., 2014; Tsuji et al., 2009). Therefore, intestinal T_regs_ play an important role in shaping cellular immune responses to mucosally encountered antigens, including oral vaccines (Gribonika et al., 2019).

Environmental Enteric Dysfunction (EED) is a gastrointestinal inflammatory disease that is most commonly seen in children in resource poor settings that contributes to developmental stunting (Keusch et al., 2014; Korpe and Petri, 2012). While EED is rare in resource rich countries, approximately 150 million children are at-risk worldwide. EED is caused by a combination of malnutrition and chronic intestinal infection/microbial dysbiosis which leads to lymphocytic infiltration, a flattening of villi in the small intestine and malabsorption due to reduced surface area (Korpe and Petri, 2012). Areas where EED is endemic are also associated with the reduced efficacy of oral vaccines (OVs) (Naylor et al., 2015). This represents a challenge, as oral vaccines are often targeted against the gastrointestinal infections that substantially impact health in resource poor settings, including contributing to the development of EED (Keusch et al., 2014). There are a variety of factors that may be contributing to the failure of oral vaccines in EED-endemic areas not the least of which is an EED-induced disruption of intestinal immunity (Bhattacharjee and Hand, 2018; Naylor et al., 2015). Using a mouse model of EED, we identify the mechanisms by which EED disrupts intestinal immune responses and the efficacy of oral vaccines. EED-affected mice exhibited a near complete block in the accumulation of intestinal CD4^+^ T cells specific to an oral vaccine but responses in secondary lymphoid tissue were unaffected. Failure in vaccine-specific CD4^+^ T cells in the intestine was accompanied by an increase in the number and function of RORγT^+^NRP1^-^ T_regs_, only in the intestine, and ablation of these cells restored oral vaccine efficacy. However, without RORγT^+^NRP1^-^ T_regs_, EED-driven stunting was significantly increased, indicating the importance of these cells in alleviating chronic intestinal inflammation. Taken together, our data supports the idea that oral vaccine failure in EED is a symptom of a localized intestinal immune regulation and uncovers a modular structure for tissue and systemic immune responses.

## Results

### Murine EED is associated with weight loss, growth stunting, and increased intestinal permeability

We sought to develop an animal model of EED which would replicate the intestinal inflammation, barrier defects and growth stunting associated with the disease in children. In previously described mouse models, changes to both the diet and microbiome are necessary to induce EED-like symptoms and in particular, affected animals showed an enrichment for *Enterobacteriaceae* in the small intestine (Brown et al., 2015; Kau et al., 2015). Therefore, we hypothesized that a combination of low protein/fat diet and colonization with an adherent-invasive *E. coli* isolate that intrinsically associated with the small intestinal epithelium would be sufficient to induce the symptoms of EED (Rolhion and Darfeuille-Michaud, 2007). The *E. coli* isolate ‘CUMT8’ was chosen because it is found naturally in mouse colonies, adheres to the small intestinal epithelium and ‘blooms’ under inflammatory conditions (Craven et al., 2012; Dogan et al., 2014). To control for diet and microbiota, our experiments had four groups: i) Isocaloric controls (ISO) fed a diet with sufficient fat and protein, ii) Malnourishment controls (MAL) fed a low protein/low fat chow, iii) CUMT8 *E. coli* colonized controls (MT8) and iv) mice which receive low protein/fat chow and are colonized with CUMT8 *E. coli* (EED) **(Fig. 1A)**. Mice were initiated to the diet at 3 weeks of age, colonized with CUMT8 *E. coli* 16, 18 and 20 days later and weighed bi-weekly throughout the protocol. All mice were fed the exact same amount of food to prevent the mice fed the low fat/protein diet from over-eating. As expected, mice fed with the malnourished diet (MAL and EED) gained weight more slowly than mice fed the isocaloric control diet (**Fig. 1B**). Independent of diet, administration of CUMT8 *E. coli* temporarily slowed the growth of mice (MT8 and EED), and the combination of low protein/fat diet and CUMT8 *E. coli* led to a 26% reduction in growth of EED mice compared to ISO controls and a significant reduction compared to MAL mice fed the same diet (**Fig. 1B and Supplemental Fig. 1A**). EED mice also exhibited other signs of growth stunting, as evidenced by reduced tail lengths (**Fig. 1C**). In humans, EED is characterized by a reduction in the length of the intestinal villi (‘blunting’) and immune infiltration (Korpe and Petri, 2012). Histological analysis of the terminal ileum revealed that EED mice, but not any of the controls exhibited villous blunting and lymphocytic infiltration in the ileum (**Fig. 1D and E**). Another symptom of EED is increased intestinal permeability (Keusch et al., 2014), and EED mice exhibited significantly increased intestinal permeability as measured by the detection orally-administered of FITC-dextran particles in the blood (**Fig. 1F**). Taken together, we have shown that this model of EED requires both a low protein/fat diet and colonization with CUMT8 *E. coli*. Conversely, gavage of CUMT8 *E. coli* alone has no obvious long-term effects on the mice fed a diet with standard amounts of fat and protein (**Fig. 1C-F**). We hypothesized that this might be due to differences in the colonization efficiency of CUMT8 *E. coli* under different diets (Brown et al., 2015). Semi-quantitative PCR analysis for the CUMT8 *E. coli* FliC gene from ileal and fecal samples revealed that MAL diet was necessary for the colonization of CUMT8 *E. coli* as mice fed a diet with sufficient protein and fat had very little CUMT8 *E. coli*, even directly following colonization (**Fig. 1G and Supplemental Fig. 1B**). In contrast, mice on the malnutrition diet had significantly (10-fold) more CUMT8 *E. coli* colonization two days after introduction compared to controls and some animals maintained colonized status for a week or more (**Fig. 1G and Supplemental Fig. 1B**). Therefore, malnourishment plays a key role in enabling adherent-invasive *E. coli* to colonize the intestine and contribute to the development of EED.

**Figure 1.**
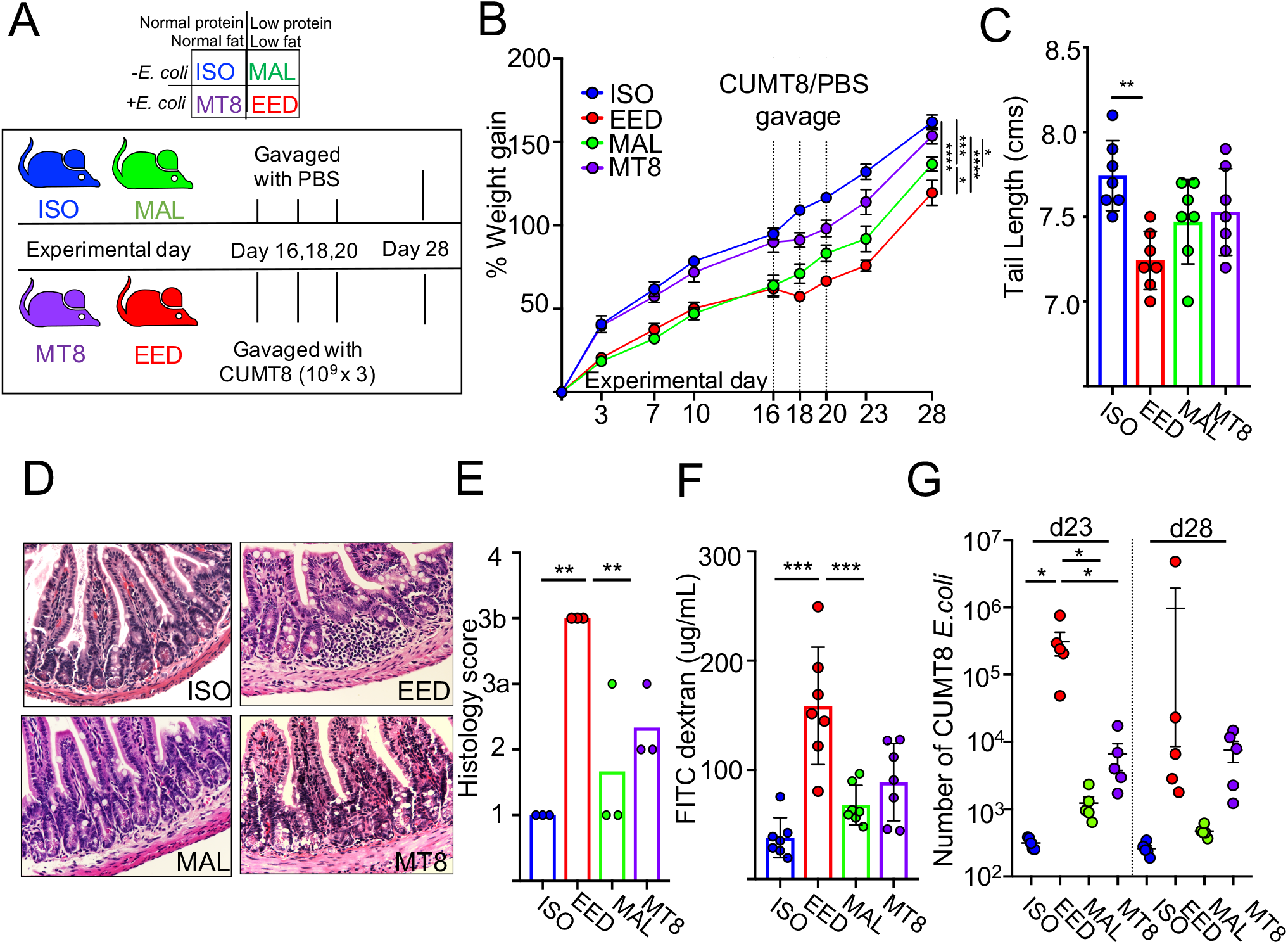
Murine EED is associated with weight loss, growth stunting and increased intestinal permeability. **A)** Schematic of the EED protocol. All mice were initiated onto the diets at weaning. ISO (blue) and MT8 (violet) mice received a control diet while MAL (green) and EED (red) mice were provided a low protein/fat diet. EED and MT8 mice were gavaged orally with 10^9^CFUs of CUMT8 *E. coli* on experimental days 16, 18 and 20. Mice were sacrificed on experimental day 28. **B)** Percent weight gain (n=4) during EED protocol. Dotted lines represent days when CUMT8 *E. coli* was orally gavaged. Data are representative of three independent experiments (n=4). **C-F** represent analysis from day 28 of the EED protocol. **C)** Tail lengths. **D)** Representative Hematoxylin and Eosin staining of ileal tissue sections(400x). **E)** Histological scoring of **D)**. **F)** Measurement of intestinal permeability determination by quantifying serum FITC Dextran (4kDA) concentration 4hrs after oral gavage. **G)** Level of CUMT8 *E. coli* in fecal samples from day 23 and day 28 of the EED protocol as calculated by semi-quantitative PCR. (**C-G**) Data points represent a single mouse. Data are represented as mean ± SEM.. (p > 0.05; *p < 0.05; **p < 0.01; ***p < 0.001). Data shown is representative of 2-7 separate experiments; **E** contains data pooled from two experiments.

### EED induces tissue-specific oral vaccine failure

Areas of the world where EED is endemic have also shown substantially reduced efficacy of oral vaccines and it has been hypothesized that EED is an important contributor to this failure (Bhattacharjee and Hand, 2018; Naylor et al., 2015). To evaluate oral vaccine responses in the murine EED model, we vaccinated mice with an attenuated Heat Labile Toxin (LT) derived from enterotoxigenic *E. coli* (ETEC). This attenuated LT, called double mutant Labile Toxin (dmLT) carries two mutations that together prevent diarrhea induction by inhibiting toxin cleavage from the plasma membrane by host proteases and reducing activation of adenylate cyclase (Clements and Norton, 2018). Despite reduced toxicity, dmLT maintains its immunogenicity as an oral vaccine and generates a robust mixed Th1/Th17 CD4^+^ T cell response and corresponding anti-LT IgA B cell response that is dependent upon immune cell activation in the draining mesenteric lymph nodes (MLNs) (Fonseca et al., 2015; Leach et al., 2012).

To test oral vaccine efficacy, at experimental day 28, mice were primed and boosted with dmLT via oral gavage (**Fig 2A**) (Hall et al., 2011). MHC class II tetramers loaded with an epitope from the A-subunit of LT (LTA_166-178_; LT-specific) were used to magnetically isolate and analyze antigen-specific CD4^+^ T cells from the small intestinal lamina propria (siLP), Peyer’s patches (PPs) and MLNs (**Fig 2B**) (Moon et al., 2009; Pepper et al., 2009). While vaccine responses were unaffected in the MLNs and PPs of EED mice, siLP anti-LT CD4^+^ T cell responses were almost undetectable in EED mice (>18 fold lower compared to ISO controls), indicating that the vaccine-specific T-cells accumulated in the secondary lymphoid tissue but not at the gut mucosa (**Fig 2B, 2C and Supplemental Fig. 2A**). EED mice also had significantly lower levels of intestinally-secreted dmLT-specific IgA compared to controls, indicating that intestinal IgA^+^ B cell responses to oral vaccination were also affected (**Fig 2D**). MAL mice also exhibited a statistically significant drop in accumulation of LT-specific CD4^+^ T cells in the siLP, suggesting that diet alone had important effects on oral vaccine efficiency but that this effect was exacerbated by EED (**Fig 2B and 2C**). Underscoring the importance of defined microbes in EED-associated oral vaccine failure, MAL mice orally gavaged with a commensal *E. coli* that does not induce EED had normal siLP LT-specific CD4^+^ T cell responses (**Supplemental Fig. 2B and C**). Taken together we have shown that in a model of EED, oral vaccines are capable of priming CD4^+^ T cell responses in the lymph nodes but that these cells fail to accumulate in the small intestine. This suggests that our murine EED model recapitulates clinical observations of substantial impairment of oral vaccine efficacy in countries where EED is common and uncovers that oral vaccine failure might be due to inhibition of intestine-resident T and B cell responses.

**Figure 2.**
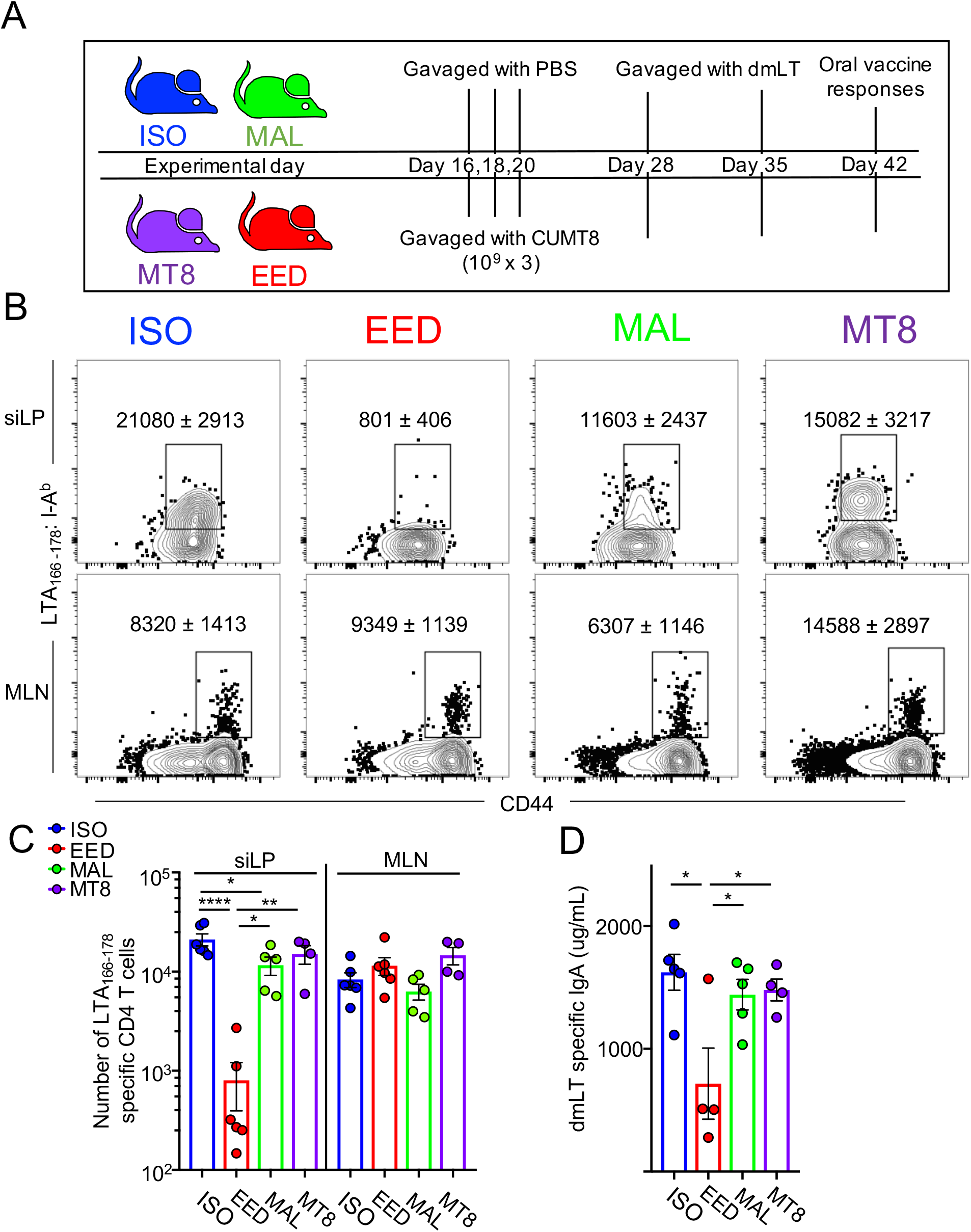
EED induces tissue-specific oral vaccine failure. **A)** Schematic of EED protocol and oral vaccination regime. Mice were immunized by oral gavage with 20μg of dmLT and boosted 7 days later. **B)** Representative flow cytometric plots of LTA_166-178_:I-A^b^ specific CD4^+^ T cells fourteen days after primary vaccination with dmLT (experimental day 42) from the siLP and MLN. Data are represented as mean ± SEM. (n=4). Cells gated: Live, Lineage^-^, CD90^+^, CD3^+^, CD8b^-^, CD4^+^. **C)** Number of LTA_166-178_:I-A^b^ specific CD4^+^ T cells calculated from **B)**. Data points represent a single mouse. **D)** Concentration of dmLT-specific IgA in the small intestine (PBS lavage). Data points represent a single mouse. **(B-D)** Data are represented as mean ± SEM. (p > 0.05; *p < 0.05; **p < 0.01; ***p < 0.001). Data shown is representative of 2-5 separate experiments; **D** shows data pooled from two experiments.

### EED-induced shifts in the microbiome are not sufficient to prevent intestinal oral vaccine responses

The intestinal microbiome of children suffering from EED has been characterized by overgrowth by *Enterobacteriaceae* and an overrepresentation of bacteria typical to the oropharyngeal cavity (Blanton et al., 2016; Kau et al., 2015; Vonaesch et al., 2018). In a healthy gut, *Enterobacteriaceae* colonization is controlled by the resident intestinal microbiota (Buffie and Pamer, 2013). We wanted to measure whether the increased efficiency of CUMT8 *E.coli* colonization in mice fed a low protein/fat diet was associated with shifts in the intestinal microbiota. To equalize the initial microbiome, mice in these experiments were purchased from a single source and randomly assorted into cages prior to the start of the protocol. Longitudinal analysis of the fecal microbiota (16S rRNA amplicon sequencing) from our four cohorts revealed that mice on the low protein/fat diet (MAL and EED) clustered away from mice fed the isocaloric diet, highlighting the effect of this diet on the microbiota (**Supplemental Fig. 3A**). As mice progressed through the protocol, principal coordinate analysis of the fecal microbiome did not significantly differentiate later fecal sampling timepoints (experimental day 23 and day 28), perhaps indicating the dominance of diet for the overall structure of the colonic microbiome in these experiments (**Supplemental Fig. 3A**). Inflammation in EED is centered on the ileum and accordingly, comparison of the ileal microbiota of ISO and EED mice at both experimental day 23 and day 28 revealed larger differences (**Supplemental Fig. 3C and D**). At experimental day 23, directly following colonization of CUMT8 *E. coli* we observed a relative increase in *Proteobacteria* (predominantly *Enterobacteriaceae*) in both ileum and fecal microbiome samples of EED mice, that was maintained (at a reduced level) until day 28 (**Supplemental Fig. 3B, D and data not shown**). Interestingly, MAL mice also showed a modest increase in fecal *Proteobacteria* at later time points, perhaps indicating the effect of this diet regimen on the outgrowth of endogenous members of this taxon (**Supplemental Fig. 3B**) (Brown et al., 2015). Therefore, in concert with clinical findings, we see that induction of EED in our murine model is associated with diet-dependent shifts in the intestinal microbiome driven by a relative expansion of *Proteobacteria*.

We next wanted to determine if EED-associated changes in the intestinal microbiome were necessary for the reduction in oral vaccine efficacy. To test this question, we treated the EED mice or control ISO mice with a broad-spectrum antibiotic cocktail (Metronidazole, Ampicillin, Neomycin and Vancomycin; MANV) to ablate the intestinal microbiome after disease has been initiated (**Fig. 3A**) (Rakoff-Nahoum et al., 2004). To limit the size of these experiments, we tested only the ISO and EED groups, which show the largest difference in oral vaccine responses. We hypothesized that ablation of the microbiota might reset local immunity and the response to oral vaccination with dmLT. However, antibiotic treatment did not significantly increase the LT-specific CD4^+^ T cell response in EED mice, which was still 10x less than ISO controls and not different from untreated EED mice (**Fig. 3B**). Thus, while the development of EED is dependent upon shifts in the intestinal microbiome, those effects on vaccine responses persist even when the EED-associated microbiome has been removed.

**Figure 3.**
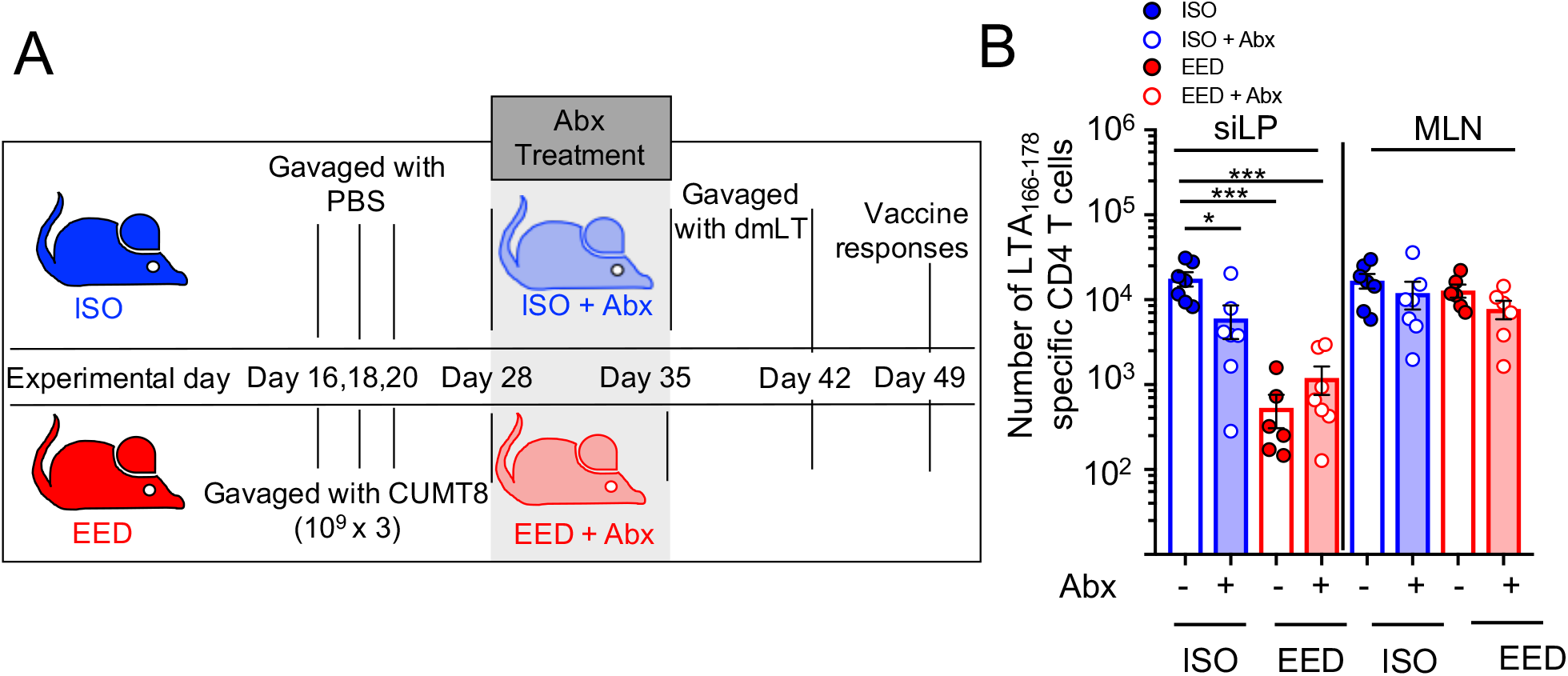
EED-induced shifts in the microbiome are not sufficient to prevent intestinal oral vaccine responses. **A**) Schematic of EED protocol, antibiotic(Abx) treatment and vaccination regime. After EED induction and prior to vaccination, mice were given a cocktail of Metronidazole, Ampicillin, Neomycin and Vancomycin (experimental day 28-35) mixed into the drinking water. Shaded area denotes duration of antibiotic treatment **B**) Numbers of LTA_166-178_:I-A^b^ specific CD4^+^ T cells isolated from the siLP and MLN of mice vaccinated after 7 days of treatment with(open circles) or without (closed circles) antibiotics. Shaded bars represent antibiotic-treated groups.Data points represent a single mouse. Data are represented as mean ± SEM. (p > 0.05; *p < 0.05; **p < 0.01; ***p < 0.001). Data shown are pooled from 2 separate experiments.

### EED-induced T_regs_ are necessary for intestinal oral vaccine failure

The intestine-specific block in oral vaccine responses in the EED mice suggested that intestinal immune microenvironment may be altered. Therefore, we evaluated the immune cells of the siLP in EED mice and controls. Previously, it was shown that infection-induced damage to the lymphatics inhibited the traffic of intestinal CD103^+^ dendritic cells (DCs) to the draining MLNs and induced large shifts in intestinal immune cell populations (Fonseca et al., 2015). Here, consistent with normal activation of LT-specific CD4^+^ T cells in the MLNs we saw no significant differences in the relative abundance of CD103^+^ DCs in the MLN and observed a modest increase of CD103^+^CD11b^+^ DCs in the siLP (**Supplemental Fig. 4A** and **B**). Since the defect in vaccine-specific CD4^+^ T cells was focused on the small intestine, we next examined major immune populations of the siLP and saw no significant differences in their proportions (**Supplemental Fig. 4C-H**). Children with EED have been reported to have increased levels of CD3^+^CD25^+^ T cells in their intestine, consistent with an increased number of FOXP3^+^ regulatory T cells (T_regs_) (Campbell et al., 2003). In accord with these findings, mice with EED were found to have significantly higher T_regs_ in their siLP, both by percent and number, compared to ISO controls (**Fig. 4A** and **B**). However, in the MLN T_reg_ percentages and numbers were invariant across all groups (**Fig. 4A** and **B**). Thus, the abundance of T_regs_ at the time of oral vaccination inversely correlated with the accumulation of LT-specific CD4^+^ T cells at that site. We next explored the possibility that the T_regs_ elicited during EED were interfering with the oral vaccine response in the gut. To test this hypothesis, EED was induced in *Foxp3^DTR-GFP^* mice, where injection with diphtheria toxin (DT) specifically and transiently ablates FOXP3^+^ T_regs_ (**Fig. 4C and Supplemental Fig. 4I**) (Kim et al., 2007). To target EED-induced T_regs_, DT was administered only for the week post-CUMT8 colonization but prior to vaccination with dmLT (**Fig. 4C**). Depletion of T_regs_ in EED mice completely restored LT-specific CD4^+^ T cell responses, while PBS-treated *Foxp3^DTR-GFP^* mice still maintained a significant impairment in these responses (**Fig. 4D and E**). As with the antibiotic depletion experiments, DT treatment was completed prior to dmLT vaccination and thus over the two week period of vaccination, T_reg_ numbers were restored to normal levels in the siLP (**Fig. 4F**), suggesting that the immune microenvironment present at the early stages of vaccination, particularly with respect to the unique EED-driven T_regs_, determines the outcome of the oral vaccine response.

**Figure 4.**
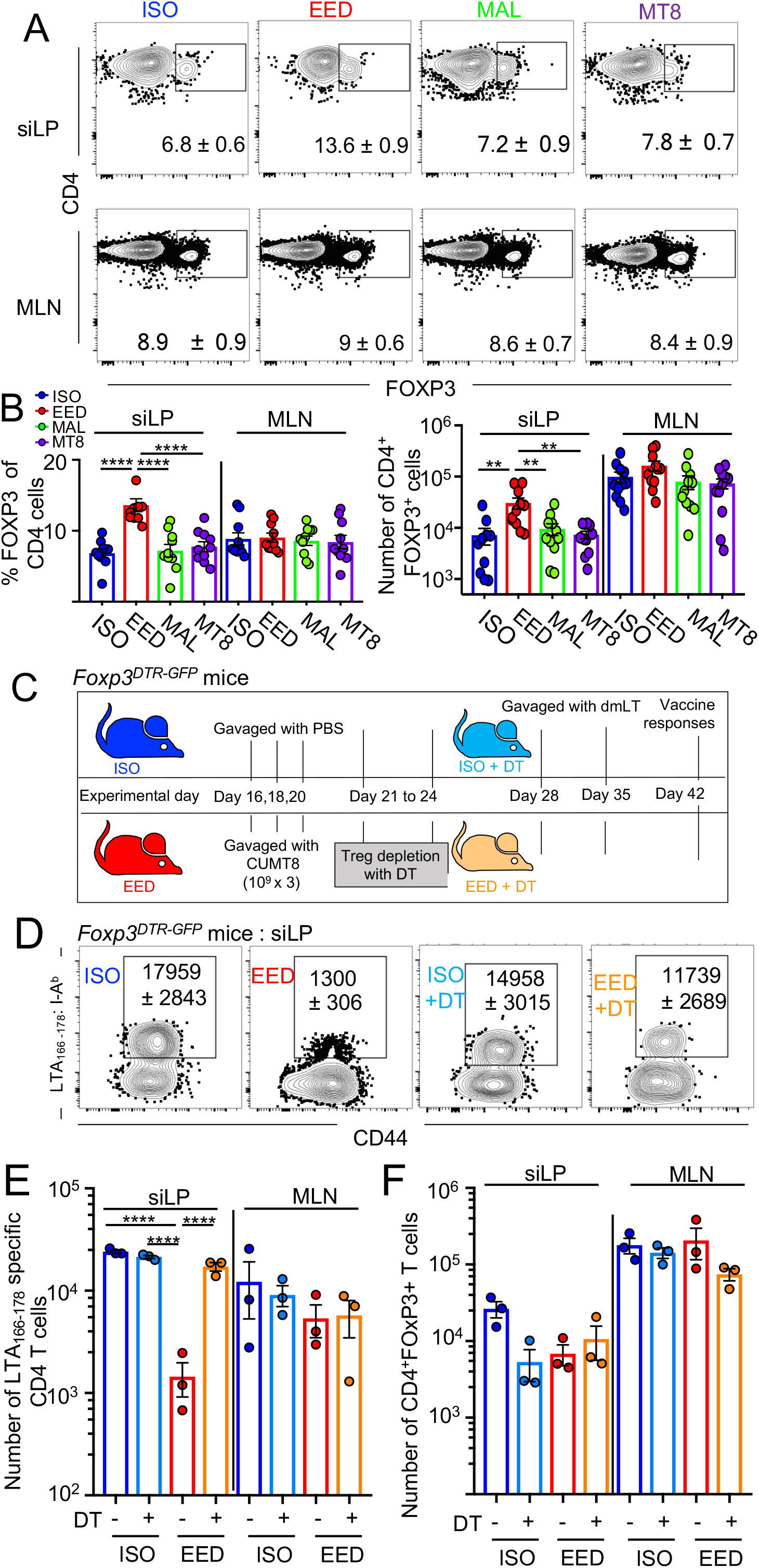
EED-induced T_regs_ are necessary for intestinal oral vaccine failure. In **A** and **B** EED is induced as in Fig.1. Lymphocytes were isolated from the small intestine at experimental day 28 of the EED protocol for analysis by flow cytometry. Gated: Live, CD90^+^, TCRβ^+^, CD8β^-^, CD4^+^ **A**) Representative flow cytometric plots showing percent FOXP3^+^ CD4^+^ T cells. **B**) Percent (left) and numbers (right) of CD4^+^FOXP3^+^ T cells from siLP and MLNs. **C**) Schematic of T_reg_ depletion experiments in mice used in **D-F**. Diphtheria toxin was injected I.P. (experimental day 21 and 24) to transiently deplete T_regs_ following gavage of CUMT8 *E. coli*. PBS was injected in controls. **D**) Representative flow cytometric plots of LTA_166-178_:I-A^b^ specific CD4^+^ T cells following oral vaccination with dmLT as in **Fig. 2**. **E**) Number of LTA_166-178_:I-A^b^ specific CD4^+^ T cells calculated from **D)**. Data points represent a single mouse. **F**) Numbers of FOXP3^+^ CD4^+^ T cells at the completion of the experiment (experimental day 42). Data points represent a single mouse. Data are represented as mean ± SEM. (p > 0.05; *p < 0.05; **p < 0.01; ***p < 0.001). Data shown is representative of 2-5 separate experiments; **B** shows data pooled from two separate experiments.

### EED induces intestinal RORγT-expressing T_regs_ that potently suppress CD4^+^ T cell responses

Given that siLP T_regs_ were necessary for the EED-mediated inhibition of oral vaccination, we wanted to determine whether EED-associated intestinal T_regs_ were functionally distinct. First, potentially accounting for the increased number of T_regs_ found in EED mice, we found that T_regs_ from EED mice expressed significantly more of the proliferation marker Ki67 than ISO mice (**Fig. 5A and B**). A critical function of T_regs_ is suppression of the clonal expansion of helper T cells. We quantified the suppression capacities of ISO and EED T_regs_ using a microsuppression assay. Strikingly, T_regs_ isolated from the siLP of EED mice had significantly greater suppressive potential than the same cells isolated from ISO controls (**Fig. 5C**). In accord with these findings, flow cytometric analysis revealed that intestinal T_regs_ from EED mice had increased expression of CD39 (**Supplemental Fig. 5A and B**), an ectonucleotidase that dephosphorylates extracellular ATP to AMP and has been associated with increased T_reg_ suppressive function (Deaglio et al., 2007). We next assessed the putative specificity (host, food or microbiota) of siLP T_regs_ by flow cytometric analysis of RORγT and NRP1 (Kim et al., 2016; Ohnmacht et al., 2015; Sefik et al., 2015; Weiss et al., 2012; Xu et al., 2018). Analysis of these markers from EED and ISO control mice revealed that siLP T_regs_ from EED mice showed a distinct increase in NRP1^-^RORγT^+^ T_regs_ (microbiota-responsive) and a corresponding loss of NRP1^+^RORγT^-^ T_regs_ (host responsive) that was specific to cells isolated from the siLP and did not extend to the draining MLNs (**Fig. 5D-F and Supplemental Fig. 5C**). Thus, we have shown that EED is associated with the specific expansion in small intestinal NRP1^-^RORγT^+^CD39^+^ T_regs_ endowed with increased regulatory function.

**Figure 5.**
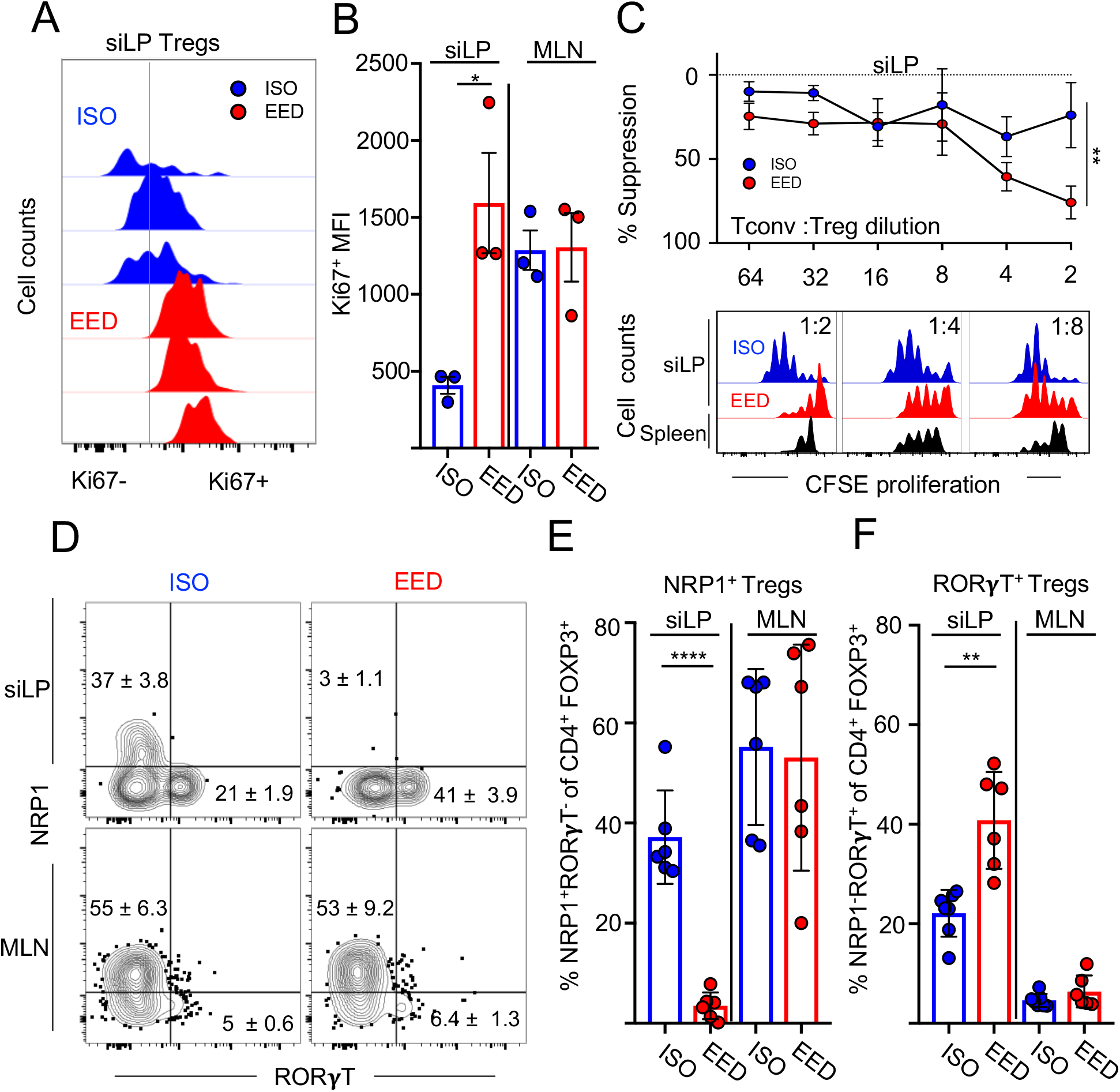
EED induces intestinal RORγT-expressing T_regs_ that potently suppress CD4^+^ T cell responses. **(A-F)** EED-like disease is induced as in Fig.1 and cells isolated at experimental day 28. **A)** Representative flow cytometric histograms showing Ki67 expression in FOXP3^+^CD4^+^ T_regs_ from the siLP. Vertical line indicates background fluorescence. **B)** Mean Fluorescence Intensity (MFI) of Ki67 expression from **A)** **C**) Percent suppression naïve CD4^+^ T cell proliferation by T_regs_ from EED mice or ISO controls (n=5). Top graph shows relative suppression at different cellular concentrations. Bottom graphs are representative histograms of T_conv_ CFSE dilution. Black histograms represent suppression by splenic T_regs_ isolated from ISO mice. Data are represented as mean ± SEM. (**p < 0.01) **D**) Representative flow cytometric contour plots of NRP1 and RORγT expression in siLP (top) and MLN (bottom) FOXP3^+^ CD4^+^ T_regs_. Gated: Live, CD90^+^, TCRβ^+^, CD8β^-^, CD4^+^, FOXP3^+^ **E)** Percent FOXP3^+^ CD4^+^ T_regs_ from **D**) that are NRP1^+^RORγT^-^. **F**) Percent FOXP3^+^ CD4^+^ T_regs_ from **D**) that are NRP1^-^RORγT^+^. **B, E and F**) Data points represent a single mouse. Bar height and error bars signify the mean and SEM (p > 0.05; *p < 0.05; **p < 0.01; ***p < 0.001). Data shown is representative of 2-3 separate experiments; **C, E** and **F** show data pooled from two experiments.

The development of NRP1^-^RORγT^+^ T_regs_ in the colon is controlled by intestinal bacteria (Abdel-Gadir et al., 2019; Sefik et al., 2015; Song et al., 2020; Xu et al., 2018). Therefore, we sought to analyze the effects of the EED microbiome on siLP T_regs_. Surprisingly, ablation of the EED-associated intestinal microbiota with antibiotics had no discernible effect on the relative abundance of NRP1^-^RORγT^+^ T_regs_ in the small intestine (**Supplemental Fig. 5D and E**). Our conclusion from these findings is that, while shifts in the EED microbiome are likely important for the development of NRP1^-^RORγT^+^ T_regs_, the microbiome is not absolutely required to sustain these cells in the small intestine. Additionally, that NRP1^-^RORγT^+^ T_regs_ are maintained independently of the microbiota may explain why antibiotic treatment failed to restore intestinal LT-specific CD4+ T cell responses (**Fig. 3B**).

### RORγT-expression in T_regs_ is necessary to inhibit oral vaccine-specific CD4^+^ T cell responses and EED-associated stunting

NRP1^-^RORγT^+^ T_regs_ have been shown to be critical to the suppression of pathology in animal models of colonic pathology (Ohnmacht et al., 2015; Sefik et al., 2015), but their role in the small intestine and specifically in oral vaccine responses has not been addressed. Therefore, we bred *Foxp3^Cre ERT2^ Rorc^fl/fl^* mice (FOXP3^RORγT+/-^) where the expression of RORγT can be specifically extinguished in FOXP3^+^ T_regs_ via the administration of tamoxifen (FOXP3^RORγT-^)(**Fig. 6A and Supplemental Fig. 6A**) (Rubtsov et al., 2010). We gavaged mice with tamoxifen to excise the *Rorc* gene in developing T_regs_ starting from five days before CUMT8 *E. coli* gavage to the end of the EED protocol. Importantly, mice treated either with tamoxifen or vehicle still developed EED, as measured by reduced weight gain throughout the experiment (**Fig. 6B**). When immunized with dmLT, tamoxifen-treated FOXP3^RORγT-^ mice showed completely restored LT-specific CD4^+^ T cell populations in the siLP, while vehicle-gavaged controls (FOXP3^RORγt+^) still showed significantly reduced responses (**Fig. 6C**). T_reg_-specific ablation of RORγT had an effect on the total number and frequency of siLP T_regs_ in EED mice, indicating that RORγT may play a role in both the accumulation and function of siLP T_regs_ (**Supplemental Fig. 6B and C**).

**Figure 6.**
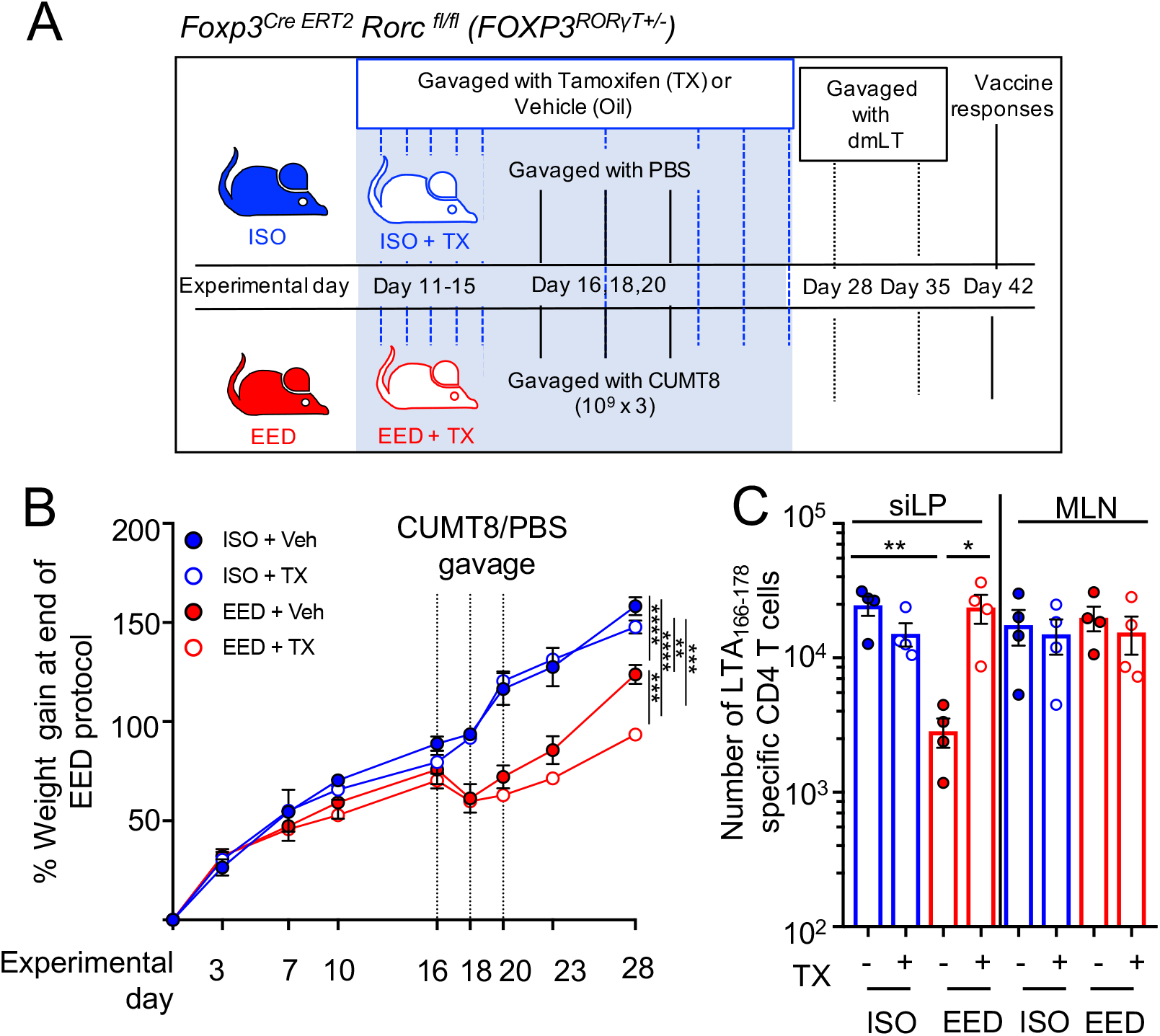
RORγT expression in T_regs_ is necessary to inhibit oral vaccine-specific CD4^+^ T cell responses and EED-associated stunting. **A)** Schematic of EED induction, tamoxifen (TX) or vehicle(Veh; corn oil) treatment, oral vaccination and RORγT deletion from FOXP3^+^ CD4^+^ T_regs_ in *Foxp3^Cre ERT2^ Rorc^fl/fl^* mice (FOXP3^RORγT+/-^). **(B-C)** Data from mice which were treated with tamoxifen(FOXP3^RORγT-^) are represented in open circles while data from vehicle treated controls (FOXP3^RORγT+^) are in closed circles. **B)** Percent weight gain (n=4) during EED establishment in *Foxp3^Cre ERT2^ RORγt^fl/fl^* mice with tamoxifen or vehicle treatment. **C)** Numbers of LTA_166-178_:I-A^b^ specific CD4^+^ T cells (in *Foxp3^CreERT2^ RORγt^fl/fl^* mice with tamoxifen or vehicle treatment isolated from the siLP and MLNs following oral vaccination with dmLT as in **Fig. 2**. Data points represent a single mouse. Data are represented as mean ± SEM. (*p < 0.05; **p < 0.01). Data shown is representative of two separate experiments; **C** shows data pooled from two experiments.

Since we treated mice with tamoxifen throughout the development of EED, we were able to determine if NRP1^-^RORγT^+^ T_regs_ are important for controlling any of the symptoms of EED. Interestingly, FOXP3^RORγt-^ EED mice were significantly more stunted than controls (FOXP3^RORγt+^) under the same protocol (**Fig. 6B and Supplemental Fig. 6D)**. Thus, RORγT expression in T_regs_ is not only necessary to inhibit the accumulation of vaccine-specific T cells in the small intestine but also important for reducing stunting associated with EED. Additionally this demonstrates the importance of NRP1^-^RORγT^+^ T_regs_ in regulating intestinal CD4^+^ T cell responses, independently of other symptoms of EED, which may be heightened by T_reg_ depletion. Altogether, we hypothesize that the accumulation of NRP1^-^RORγT^+^ T_regs_ in the intestine of EED may result from an effort to alleviate chronic intestinal inflammation in the small intestine and that the localized failure of vaccine-specific T cell responses at this site develops as a side effect of a tissue-specific immunoregulation.

## Discussion

Here we have shown that a disease that mimics several aspects of human EED can be induced in mice with the combination of a low protein/fat diet and colonization with adherent invasive *E. coli*. Areas of the world where EED is endemic also exhibit a lack of efficacy of oral vaccines, which is tragic since these communities would probably benefit most from effective mucosal immune protection (Grassly et al., 2009; Levine, 2010). There are a variety of factors that contribute to the failure of oral vaccines in low-to-middle income countries (LMICs), including extended breast feeding, which may contain vaccine inhibitory IgA and milk oligosaccharides, increased exposure to pollution, micronutrient deficiency, pathogen heterogeneity and EED (Bhattacharjee and Hand, 2018; Moon et al., 2010; Santos and Hoshino, 2005). Here, we show that independent of other factors, EED is sufficient to inhibit siLP CD4^+^ T cell responses and IgA production, adding evidence for the hypothesis that EED is partially responsible for the failure of oral vaccines. Further, we identify small intestinal T_regs_ as the mechanism of the inhibition of oral vaccine responses, which will focus future studies and therapies.

Most research on oral vaccines has focused on antibodies. However, there is a growing interest in the potential of intestinal-resident CD4^+^ T cells in mediating protection via the rapid production of cytokines such as IL-17A and IL-22, that can activate defense mechanisms in the intestinal epithelium (Cao et al., 2012; Omenetti et al., 2019; Schreiber et al., 2015). A recent trial using oral dmLT in combination with live-attenuated enterotoxigenic *E. coli* vaccine in Bangladesh revealed significant protection that was not correlated to antigen-specific IgA or IgG, leaving the possibility of a T cell-mediated mechanism (Clements and Norton, 2018; Harro et al., 2019). CD4^+^ T cells also support the differentiation and development of IgA^+^ plasma B cells in Peyer’s Patches (PP) (Bemark et al., 2016; Lycke and Bemark, 2017), but, surprisingly, we saw very few LT-specific T cells in the PP in our studies and they did not vary according to nutritional/EED status. After activation in the PP, IgA^+^ plasma cells traffic to siLP where they will reside long-term (Lycke and Bemark, 2017). Perhaps siLP CD4^+^ T cells assist B cells in the small intestine or alternatively, the inflammatory and nutrient-deficient environment of the EED intestine may inhibit IgA-producing B cells independently of T cell help. Future studies on LT-specific B cells in EED will be necessary to elucidate this point.

In our model of EED, we correlated oral vaccine failure with an increased presence of NRP1^-^ RORγT^+^ T_regs_, within the small intestine but not in the draining MLN tissue. This is consistent with human studies where children suffering from EED have increased CD3^+^CD25^+^ cells in their intestine (Campbell et al., 2003). Most previous descriptions of NRP1^-^RORγT^+^ T_regs_ have been focused on the colon, where these cells are dependent upon the microbiota and its production of secondary bile acid metabolites (Sefik et al., 2015; Song et al., 2020; Xu et al., 2018). Only a fraction (5%) of bile acids end up in the colon and the vast majority are recycled by the ileum (Ridlon et al., 2006), so it is somewhat dissonant that NRP1^-^RORγT^+^ T_regs_ would be more common in the colon than ileum. The type of metabolites produced by the microbiome may differ between the small intestine and colon, which could account for this difference. Further, the metabolite profile of the small intestine is substantially different between EED mice and controls with bile acid metabolism/synthesis amongst the largest differences (Brown et al., 2015). Alternatively, NRP1^-^RORγT^+^ T_regs_ have been observed in lung tissue, where bile acids are not present, implying that there may be multiple pathways that drive this subset (Lochner et al., 2008). The cytokines IL-6, IL-10 and IL-23 are also important for the induction and maintenance of NRP1^-^RORγT^+^ T_regs_ and perhaps intestinal damage during EED induces these cytokines in the small intestine (Kim et al., 2018; Ohnmacht et al., 2015). In addition, while the microbiome was clearly important for the development of NRP1^-^RORγT^+^ T_regs_, ablation of the microbiome with antibiotics after their development had no measurable effect on their presence in the small intestine over the period we observed. This might represent a key difference in the signals required for persistence of NRP1^-^RORγT^+^ T_regs_ in different parts of the intestine. Alternatively, antigen-specific NRP1^-^RORγT^+^ T_regs_ are induced by colonization with *Helicobacter hepaticus* (Xu et al., 2018). It is not clear whether the NRP1^-^RORγT^+^ T_regs_ induced by EED are specific to CUMT8 *E. coli* but both *H. hepaticus* and CUMT8 *E. coli* share the ability to adhere to the surface of the intestine (Dogan et al., 2014; Fox et al., 1994). Adherence to the gut lining, is likely critical for the induction of NRP1^-^RORγT^+^ T_regs_ (Kim et al., 2018), as seen by the lack of oral vaccine failure in malnourished mice that were given ECMB, a non-adherent strain of *E.coli*. Regardless of their antigen-specificity, it will be interesting to determine if NRP1^-^RORγt^+^ T_regs_ have memory functions that allow them to survive without constant signals from the microbiome. Critically, patients suffering from EED can take years to re-establish full intestinal functionality, and it will be important to determine whether, once established, intestinal resident immune cells maintain the disease state long after the inciting conditions have subsided (Gerson et al., 1971; Lindenbaum et al., 1972).

The molecular mechanism by which NRP1^-^RORγT^+^ T_regs_ limit the accumulation of LT-specific CD4^+^ T cells is not clear. Our hypothesis is that NRP1^-^RORγT^+^ T_regs_ are blocking local antigen presentation and costimulation that is necessary for secondary proliferation once LT-specific T cells arrive in the intestine (Sano et al., 2015). The function that RORγT performs as a transcription factor in T_regs_ is also not clear at this point. Our results showed that RORγT expression may be important for T_reg_ proliferation and survival in the intestine. Loss of RORγT expression from T_regs_ does not lead to spontaneous intestinal inflammation but, instead, is critical to limiting inflammation once it is induced (Boehm et al., 2012; Ohnmacht et al., 2015; Sefik et al., 2015). Accordingly, RORγT^+^ T_regs_ were superior to RORγT^-^ T_regs_ at suppressing inflammation in a model of T cell transfer colitis(Yang et al., 2016). In concert, our results show that NRP1^-^RORγt^+^ T_regs_ reduce EED-associated stunting. Since the primary function of the gut is to absorb nutrients, we hypothesize that NRP1^-^RORγt^+^ T_regs_ prioritize epithelial health and absorption and that the block in immune responses is a by-product of the necessity to regulate damaging inflammation.

Adherent-Invasive *E. coli* were first described in patients with Crohn’s Disease where it contributes substantially to pathology, but are also present amongst the general population (Barnich and Darfeuille-Michaud, 2007). It is likely that our ancestors were commonly colonized with bacteria that irritated the epithelium and engendered CD4^+^ T helper responses. We propose that EED and augmented intestinal adherence of *Enterobacteriaceae*, is inducing a local immune suppression, mediated by NRP1^-^RORγt^+^ T_regs_, to limit inflammatory intestinal damage. In this model, systemic immunity is left intact to guard against more invasive organisms. However, a side effect of this EED-induced intestinal immune shutdown is the loss of efficient tissue responses to oral vaccines. We hypothesize that this represents an evolutionary mechanism to prevent chronic inflammation in the small intestine, such as Inflammatory Bowel Disease in the face of constant colonization with intestinally-adherent bacteria. In the future, as we develop deeper knowledge of the mechanisms underpinning the intestinal immune defect in EED, we hope that these will be incorporated into treatment plans and oral vaccine formulations so we can break the chain of malnutrition and infection that underlies this disease.

## Acknowledgments

This work was supported by the National Institutes of Health (R21AI142051; TWH, NIAID division of Intramural Research; YB), the R.K. Mellon Institute for Pediatric Research (TWH), and the UPMC Children’s Hospital of Pittsburgh RAC award (AB). AEO is supported by a Damon Runyon Cancer Fellowship and AHPB is supported by an NIH T32 (T32AI089443). The authors would like to thank J. Michel and A. Styche for assistance with cell sorting, the Children’s Hospital of Pittsburgh Histology core for the preparation of tissue slides, and the University of Pittsburgh Division of Laboratory Animal Research. We would also like to thank D. Vignali (Foxp3^eGFP-Cre-ERT2^), K. Simpson (CUMT8 *E. coli*) and J. Williams (Foxp3^DTR-GFP^) for sharing critical reagents and M. Pepper and M. Jenkins for assistance with making and using MHC class II tetramer reagents. We would like to thank the University of Pittsburgh Center for Research Computing for resources and assistance with genetic sequence analysis. We would like to thank the members of the Hand, Belkaid, Poholek and Canna labs for discussion and critical reading of the manuscript.

## Author Contributions

AB, AEO and TWH designed the experiments; AB, AHPB, AEO, JTT, DY and BRH carried out the experiments; AB, AHPB, AEO and TWH analyzed the data; JLL, SPS, JAH, OJH, DMF and YB contributed to the development of reagents (MHC II tetramers) and models to study anti-LT responses in vivo; EBN provides recombinant LT and dmLT for these studies; AB and TWH wrote the manuscript.

## Materials and Methods

### Mice

3-week old wild type C57BL/6Tac mice (B6 MPF; Taconic) were used for EED establishment and characterization unless otherwise noted. Foxp3^DTR-GFP^ (*Foxp3^tm3(DTR/GFP)Ayr^*) mice, were the kind gift of John Williams (UPMC Children’s Hospital of Pittsburgh) (Kim et al., 2007). Foxp3^eGFP-Cre-ERT2^ (*Foxp3 ^tm9(EGFP/cre/ERT2)Ayr^*) mice were the kind gift of Dario Vignali (University of Pittsburgh) and bred with Rorc^fl/fl^ mice (B6(Cg)-^Rorctm3Litt^) from Jackson Laboratories (Choi et al., 2016; Rubtsov et al., 2010). Both male and female age-matched mice (3 weeks) were used for all experiments. All experiments were performed in an American Association for the Accreditation of Laboratory Animal Care-accredited animal facility at the University of Pittsburgh. Mice were kept in specific pathogen-free conditions and housed in accordance with the procedures outlined in the Guide for the Care and Use of Laboratory Animals under an animal study proposal approved by the Institutional Animal Care and Use Committee of the University of Pittsburgh.

### Bacterial strains

CUMT8, an autochthonous mouse adherent invasive *E. coli* (Dogan et al., 2014), was provided by Kenneth Simpson (Cornell University, Ithaca, NY). In some experiments, a mouse commensal *E. coli* (ECMB) was used in place of CUMT8 (Molloy et al., 2013). ECMB lacks a flagellar operon and is non-motile.

### EED Induction

Immediately following arrival, mice are put on one of two special diets (Research Diets, New Brunswick, NJ); a malnourished low protein/low fat diet (7% protein/7% fat; D14071001)), or a control isocaloric diet (20% protein/15% fat; D09051102) (Brown et al., 2015). Chow was added bi-weekly to ensure that malnourished mice did not compensate protein/fat deficiencies by increasing intake and all mice were provided the same amount of calories each week. After 2 weeks on defined diets, some mice (MT8 and EED) were orally gavaged with 1×10^9^ CUMT8 *E. coli* on experimental days 16, 18 and 20 post-weaning. Mice were weighed weekly. After 28 days on diets, mice were sacrificed and evaluated for signs of EED. Alternatively, mice were then treated or immunized at experimental day 28.

### Histological analysis of terminal ileums

Terminal ileum samples were fixed in formalin, dehydrated and paraffin embedded. Sections of ileum were stained with hematoxylin and eosin stains for morphological analysis and slides were scored using the modified Marsh-Oberhuber classification (Dickson et al., 2006). Outline of scoring system:

0=Pre-infiltrative, 1=Infiltrative, 2=Infiltrative-hyperplastic, 3a=Flat destructive with mild villous atrophy, 3b= Flat destructive with moderate villous atrophy, 3c= Flat destructive with total villous atrophy and 4=Atrophic-hypoplastic.

### FITC–dextran assay

For evaluation of gut permeability, mice were orally gavaged with 4 kDa FITC–dextran (Sigma-Aldrich) after fasting for 4 hrs. FITC–dextran (100 mg/ml) was dissolved in PBS and gavaged at 44mg/100g body weight. Four hours post gavage, mice were euthanized and blood was collected immediately via cardiac puncture. Serum was isolated and was diluted with an equal volume of PBS, of which 100μl which was added to a 96-well microplate in duplicate. FITC in blood sera was quantified using florescence spectroscopy. The plate was read at an excitation of 485 nm (20 nm band width) and an emission wavelength of 528 nm, and serially diluted FITC-dextran was used to calculate serum concentrations. A naïve mouse gavaged with PBS for the same duration was used to determine background fluorescence.

### Oral Vaccination

Double mutant *E. coli* heat labile toxin (R192G/L211A) (dmLT), was produced from *E. coli* clones expressing recombinant protein as previously described (Norton et al., 2011). Mice were immunized twice, 7 days apart by oral gavage and vaccine responses were assayed 2 weeks after primary gavage as described before (Hall et al., 2011).

### Antibiotic treatment

The antibiotic cocktail used comprised of Metronidazole (1mg/mL), Ampicillin (1mg/mL), Neomycin (1mg/mL) and Vancomycin (0.5mg/mL) (MANV). Antibiotics were provided in the drinking water and were given to mice for one week after EED establishment (days 28 to 35 on diets). To reduce the bitter taste of metronidazole and prevent dehydration, antibiotic water was supplemented with 10g/L of saccharin (Sweet ‘N Low).

### Real-time qPCR analysis for bacterial abundance

DNA from stool and ileal contents were isolated using QIAamp Fast DNA Stool Mini Kit (Qiagen) followed by real-time RT-PCR (qPCR) using SYBR Green supermix (Bio-RAD) on a CFX Connect Real-time PCR Detection Machine (Bio-RAD). CUMT8 *E. coli* was measured using primers (Fwd 5’ CGACAGTGCCGGTATTTGTA, Rev 5’ TGAGTACTGCGGATGGTTCA) that amplify the FliC gene and number of bacteria calculated against a standard curve prepared from known quantities of CUMT8 *E. coli*.

### 16S rRNA amplicon analysis of relative bacterial abundance

DNA was isolated using the QIAamp Fast DNA Stool Mini Kit (Qiagen) and the extracted DNA was stored at −20 °C prior to 16S amplicon PCR and sequencing. Analysis of small subunit ribosomal RNA gene (16S rRNA) expression was done using following primers;16S primer Fwd: 5’-ACTCCTACGGGAGGCAGCAGT-3’, Rev: 5’-ATTACCGCGGCTGCTGGC-3’. Sequencing was carried out by BGI Genomics. Microbiome informatics were performed using QIIME2 2020.2 (Bolyen et al., 2019). Raw sequences were quality-filtered and denoised with DADA2 (Callahan et al., 2016). Amplicon variant sequences (ASVs) were aligned with MAFFT and used to construct a phylogeny with FastTree2 (Katoh et al., 2002). Alpha diversity metrics (observed OTUs), beta diversity metrics (Bray Curtis dissimilarity) and Principle Coordinate Analysis (PCA) were estimated after samples were rarefied to 63,000 (subsampled without replacement) sequences per samples. Clustering of fecal and ileal microbiota samples was performed by PCA based on bacterial community similarity (using the Bray Curtis algorithm), with ellipsoids representing 95% CI for each group assuming multivariate t-distribution. Taxonomy was assigned to ASVs using naive Bayes taxonomy classifier against the Greengenes 18_8 99% OTUs reference sequences (McDonald et al., 2012). All plots were made with publicly available R packages.

### Flow cytometry

All antibodies used for flow cytometry were purchased from either ThermoFisher, BD Biosciences, or BioLegend. The flowing antibodies were used : CD45(104), CD90(30-H12), CD4(RM4-5), CD8b(H35-17.2), CD44(IM7), FOXP3(MF23), NRP1(3E12), RORγT(B2D), CD11b(M1/70), CD11c(N418), Ly6G(1A8), SiglecF (IRNM44N), Ly6C(Hk1.4), MHCII(M5/114), CD64(X54-5/7.1), CD39(Duha59), CD73(TY/11.8), TCRβ (H57-597), Ki67(16A8), CD3(17A2), CD19(1D3). Dead cells were discriminated in all experiments using LIVE/DEAD fixable dead stain (ThermoFisher). All stains were carried out in media containing anti-CD16/32 blocking antibody (clone 93, ThermoFisher). For intracellular staining, cells were fixed and permabilized using the FoxP3/Transcription factor staining buffer set according to the manufacturer’s directions (ThermoFisher). APC-conjugated MHC class II (I-A^b^) LT_166-178_ tetramers (RYYRNLNIAPAED) were produced in our laboratory as previously described (Moon et al., 2007). Staining of tetramer positive T cells was carried out after magnetic isolation of the cells as described (Moon et al., 2009). All tetramer stains were performed at room temperature for 45–60 minutes. All flow cytometry was acquired on an LSRFortessa FACS analyzer and cell sorting was carried out on a FACS Aria (BD Biosciences). Cells were isolated from the small intestine, Peyer’s Patches, spleen and lymph nodes for flow cytometry as described previously (Hand et al., 2012).

### Suppression assay

Small intestinal T_regs_ (CD45^+^CD90^+^CD4^+^GFP^+^ T cells) from ISO and EED *Foxp3^DTR-GFP^* mice were sorted using the BD FACSAria sorter for comparing suppressive efficacy along with splenic non-T_regs_(CD45^+^CD90^+^CD4^+^GFP^-^ T cells) cells and splenic antigen presenting cells (APCs) (CD45^+^CD90^-^) from a naïve mouse. The responder cells (splenic non-T_regs_) from naïve *Foxp3^DTR-GFP^* mice were labeled with 5μM CellTrace Violet (Life Technology). Responder cells (4×10^3^), APCs (8×10^3^), and different concentrations of T_regs_ (1:2-1:64 T_reg_: T_eff_ ratio, 500-2000 T_regs_) were activated with 2μg/ml anti-CD3 (Biolegend) in a 96-well round bottom plate with 100ul RPMI for 72hrs (Turnis et al., 2016). Suppression was calculated as previously described (McMurchy and Levings, 2012). Briefly, cells were acquired by the BD LSR Fortessa™, and the division index of responder cells was analyzed on the division of CellTrace Violet. Suppression was then calculated with the formula % Suppression = (1-DI_Treg_/DI_Ctrl_) × 100% (DI_Treg_ stands for the division index of responder cells with T_regs_, and DI_Ctrl_ stands for the division index of responder cells activated without T_regs_).

### Anti-dmLT ELISA

For detection of LT-specific IgA antibody from intestinal lavage, 96-well-plates were coated with 1μg/ml of dmLT protein for overnight at 4°C, blocked with 1% BSA for 1 hour followed by incubation with intestinal lavage (cleared of solid particles/bacteria by centrifugation) overnight at 4°C. Next day, after washing, IgA antibody bound to dmLT was detected by HRP-labeled anti-IgA antibody and developed using standard ELISA substrate. A naïve mouse was used as a negative control.

### Diphtheria toxin (DT) and Tamoxifen (TX) administration

Mice were intraperitoneally (I.P.) injected with 25ug/kg diphtheria toxin (DT, Sigma) twice, with 3 days between injections (PBS used as vehicle). Tamoxifen (TX, Sigma-Aldrich), was orally gavaged at 5mg/mouse/day, first for 5 consecutive days to activate the promoter and then once every 3 days for the duration of the deletion. Since TX is poorly soluble in water, so the amount needed for a single day was dissolved in EtOH with heating to 37°C and then diluted in corn oil (Sigma) such that 100ul had 5mg. Corn oil was used for vehicle controls.

### Statistics

Due to the necessities of diets and microbiome modification, mouse studies were performed in a non-blinded manner. Histology was scored blinded. Data are presented as mean ± SEM. Statistical significance was determined using unpaired Student’s t test when comparing two groups, and one-way ANOVA with multiple comparisons, when comparing multiple groups. Experiments comparing the weight of animals over time were evaluated with a two-way ANOVA with multiple comparisons; row means were analyzed. All statistical analysis was calculated using Prism software (GraphPad). For details on significance, please see figure legends.

## Supplemental Figure Legends

**Supplemental Figure 1 (Related to Fig.1).**
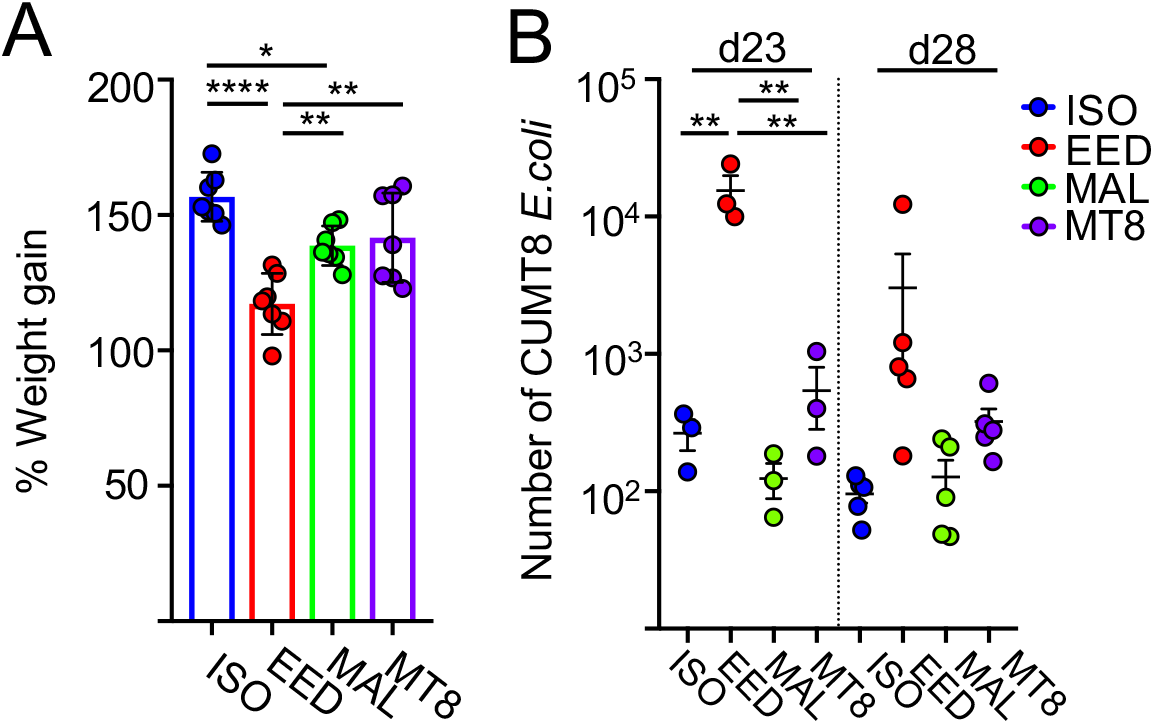
**A)** Percent weight gain at experimental day 28 of the EED protocol. **B)** Level of CUMT8 *E. coli* in ileal samples from experimental day 23 and day 28 of the EED protocol as calculated by semi-quantitative PCR. Data points represent a single mouse. Data are represented as mean ± SEM. (p > 0.05; *p < 0.05; **p < 0.01; ***p < 0.001). Data shown is representative of 2-7 separate experiments.

**Supplemental Figure 2 (related to Figure 2).**
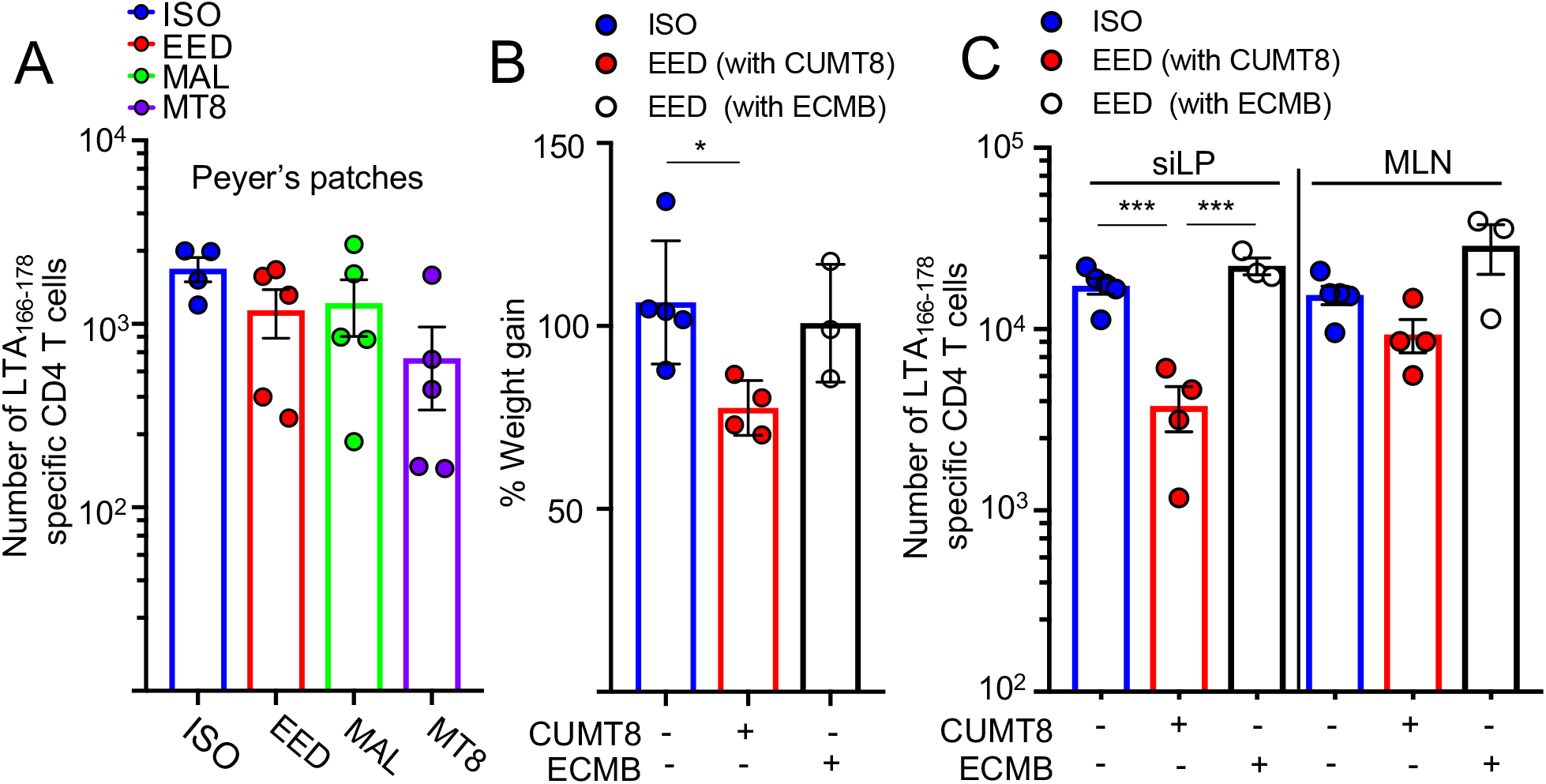
**A)** Number of LTA_166-178_:I-A^b^ specific CD4^+^ T cells isolated from Peyer s Patches. **B** and **C** Comparison of ISO, EED and a third group of mice fed the low protein/fat chow but gavaged with a commensal *E. coli* ‘MB’ (ECMB). **B**) Percent weight gain at experimental day 28 of the EED protocol. **C**) Numbers of LTA_166-178_:I-A^b^ specific CD4^+^ T cells in the siLP and MLN. Data points represent a single mouse. Data are represented as mean ± SEM. (p > 0.05; *p < 0.05; **p < 0.01; ***p < 0.001).

**Supplemental Figure 3 (related to Figure 3).**
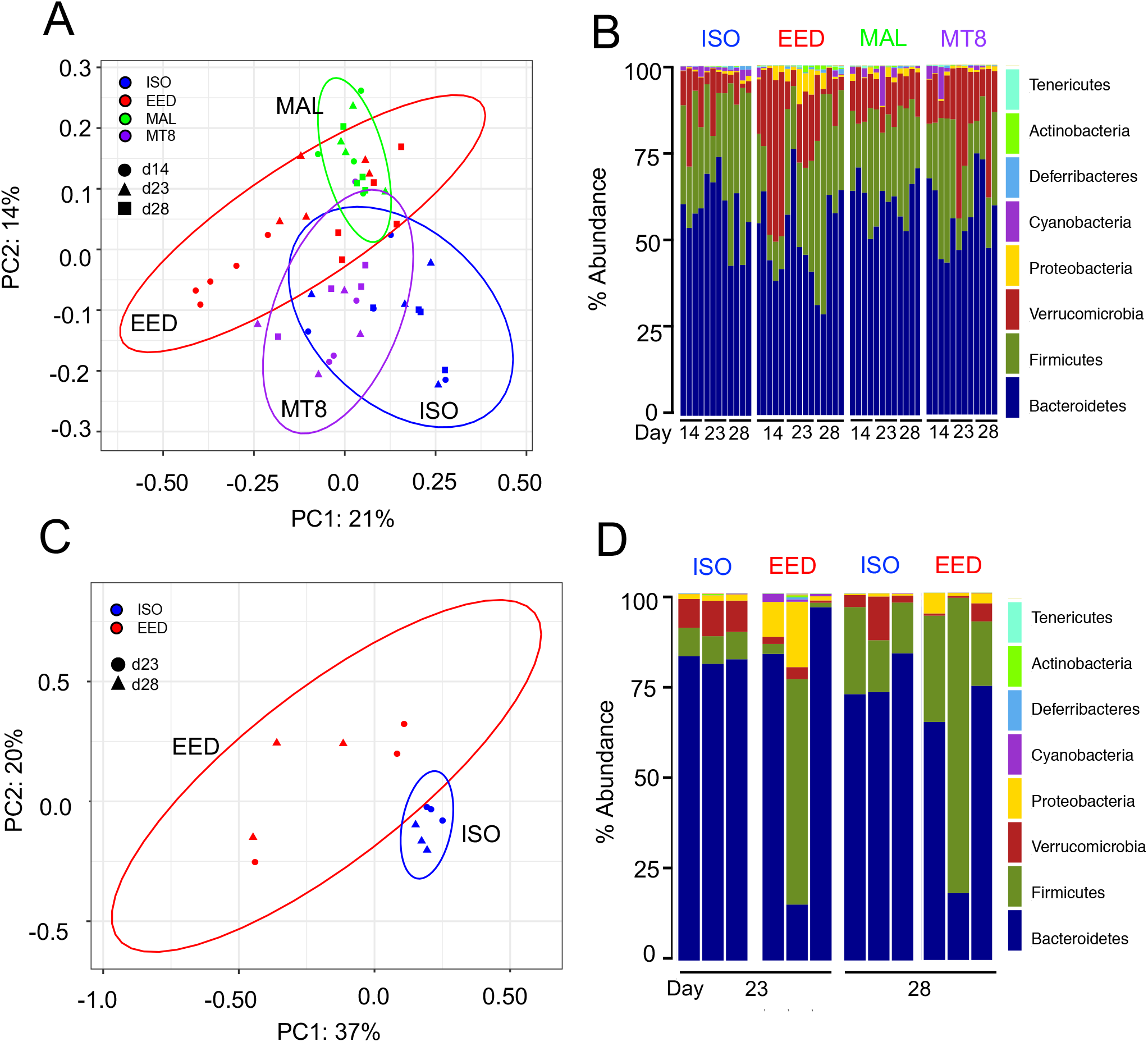
**A-D** Genomic DNA was isolated from fecal and ileal samples and the structure of the microbiome analyzed via the expression of 16S rRNA genes. **A**) PCA of fecal samples collected on experimental day 14, 23 and 28 based on bacterial community similarity, with ellipsoids representing 95% CI for each group assuming multivariate t-distribution. **B**) The mean relative abundances of the most abundant phylum level OTUs in fecal samples (n=4-5) on experimental day 14, 23 and 28. **C**) PCA of ileal samples based on bacterial community similarity, with ellipsoids representing 95% CI for each group assuming multivariate t-distribution. **D)** The mean relative abundances of the most abundant phyla in ileal samples isolated on experimental day 23 and 28. Data shown is representative of two separate experiments.

**Supplemental Figure 4 (related to Fig. 4).**
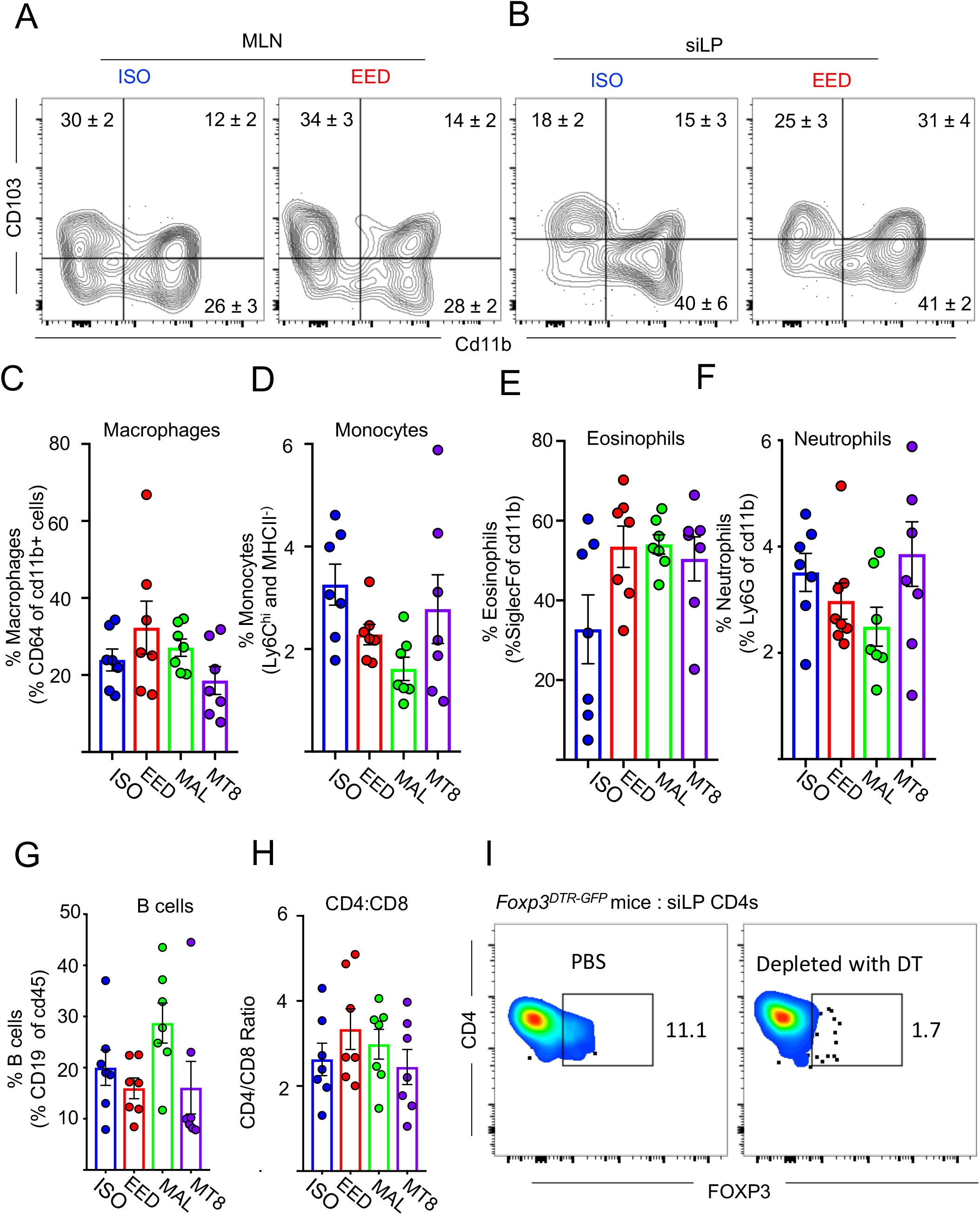
**A**) Representative flow cytometric contour plots of dendritic cell subsets from the MLN (Gated on Live CD45^+^ Ly6G^-^ SiglecF^-^ Ly6C^-^ CD90^-^ CD19^-^ CD64^-^ CD11c^+^ MHCII^+^) **B**) Representative flow cytometric contour plots of dendritic cell subsets from the siLP (Gated on Live CD45^+^ Ly6G^-^ SiglecF^-^ Ly6C^-^ CD90^-^ CD19^-^ CD64^-^ CD11c^+^ MHCII^+^). For **A** and **B** numbers denote the mean±SEM (n=3). (**C-H**) Immune cell phenotyping of siLP resident immune cells. **C**) Relative frequency of macrophages (Gated Live CD45^+^ Ly6G^-^ SiglecF^-^ Ly6C^-^ CD90^-^ CD19^-^ CD64^+^ CD11b^+^) **D**) Relative frequency of monocytes (Gated Live CD45^+^ Ly6G^-^ SiglecF^-^ CD90^-^ CD19^-^ CD64^-^ CD11c^-^ MHCII^-^ Ly6C^+^ CD11b^+^) **E**) Relative frequency of eosinophils (Gated Live CD45^+^ Ly6G^dim^ SiglecF^+^ CD90^-^ CD19^-^ CD64^-^CD11 c^-^ MHCII^-^ Ly6C^-^ CD11 b^+^) **F**) Relative frequency of neutrophils (Gated Live CD45^+^ Ly6G^+^ SiglecF^-^ CD90^-^ CD19^-^ CD64^-^CD11 c^-^ MHCII^-^ Ly6C^+^ CD11 b^+^) **G**) Relative frequency of B cells (Gated Live CD45^+^ Ly6G^-^ SiglecF^-^ CD90^-^ CD19^+^) **H**) Ratio of CD4^+^ to CD8β^+^ T cells (T cells gated Live CD45^+^ CD90^+^ TCRβ^+^ CD4^+/-^ CD8β^+/-^ **I**) Representative flow cytometric plot showing depletion of CD4^+^ FOXP3^+^ cells from the siLP following diptheria toxin administration as described in **Fig. 4C**(n=3). Data points represent a single mouse. Data are represented as mean ± SEM. (p > 0.05; *p % 0.05; **p % 0.01; ***p % 0.001). Data shown is representative of 3 separate experiments; **C-H** shows data from two pooled experiments.

**Supplemental Figure 5 (related to figure 5).**
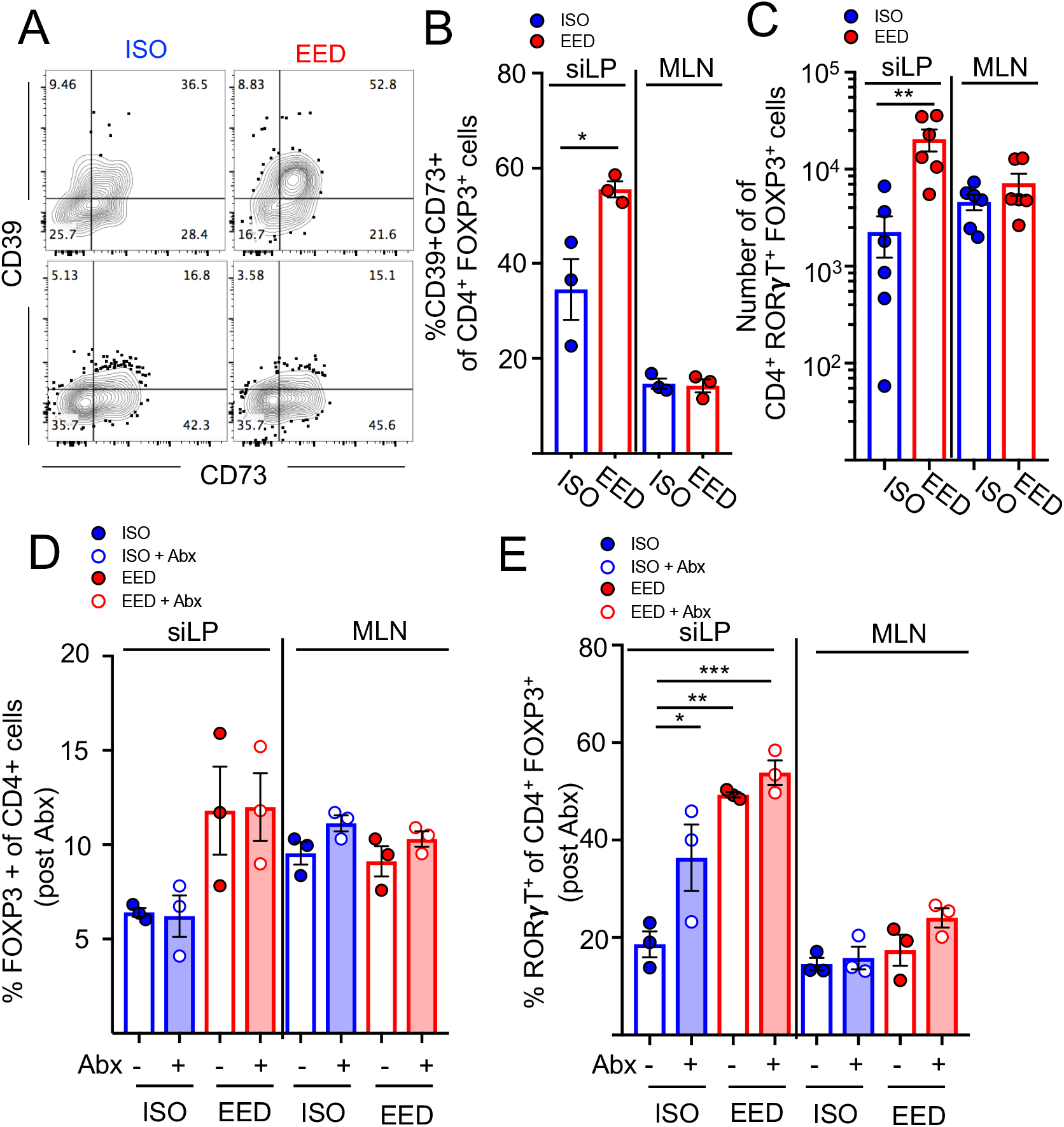
**(A-C)** Experimental day 28 of the EED protocol. **A)** Representative flow cytometric contour plots of siLP (top) and MLN (bottom) FOXP3^+^ CD4^+^ T_regs_. **B**) Percent of CD39^+^ CD73^+^ of FOXP3^+^ CD4^+^ T_regs_ from **A**). **C**) Numbers of RORγT^+^ FOXP3^+^ CD4^+^ T_regs_. **(D and E)** Mice were treated with antibiotics (MANV-see Fig 3A or methods for details of treatment). Shaded bars represent antibiotic-treated groups. **D**) Percent FOXP3^+^ CD4^+^ T_regs_ (gated on CD4s) in the siLP and MLN of mice treated with (open circles) or without (closed circles) 7 days of antibiotics, as in **Fig. 3**. **E**) Percent RORγT^+^ cells of FOXP3^+^ CD4^+^ T_regs_ in the siLP and MLN of mice mice treated with (open circles) or without (closed circles) 7 days of antibiotics, as in **Fig. 3**. Data points represent a single mouse. Data are represented as mean ±SEM. (p > 0.05; *p < 0.05; **p < 0.01). Data shown is representative of 1-2 experiments; **C** shows pooled data from two experiments.

**Supplemental Figure 6 (related to Fig. 6).**
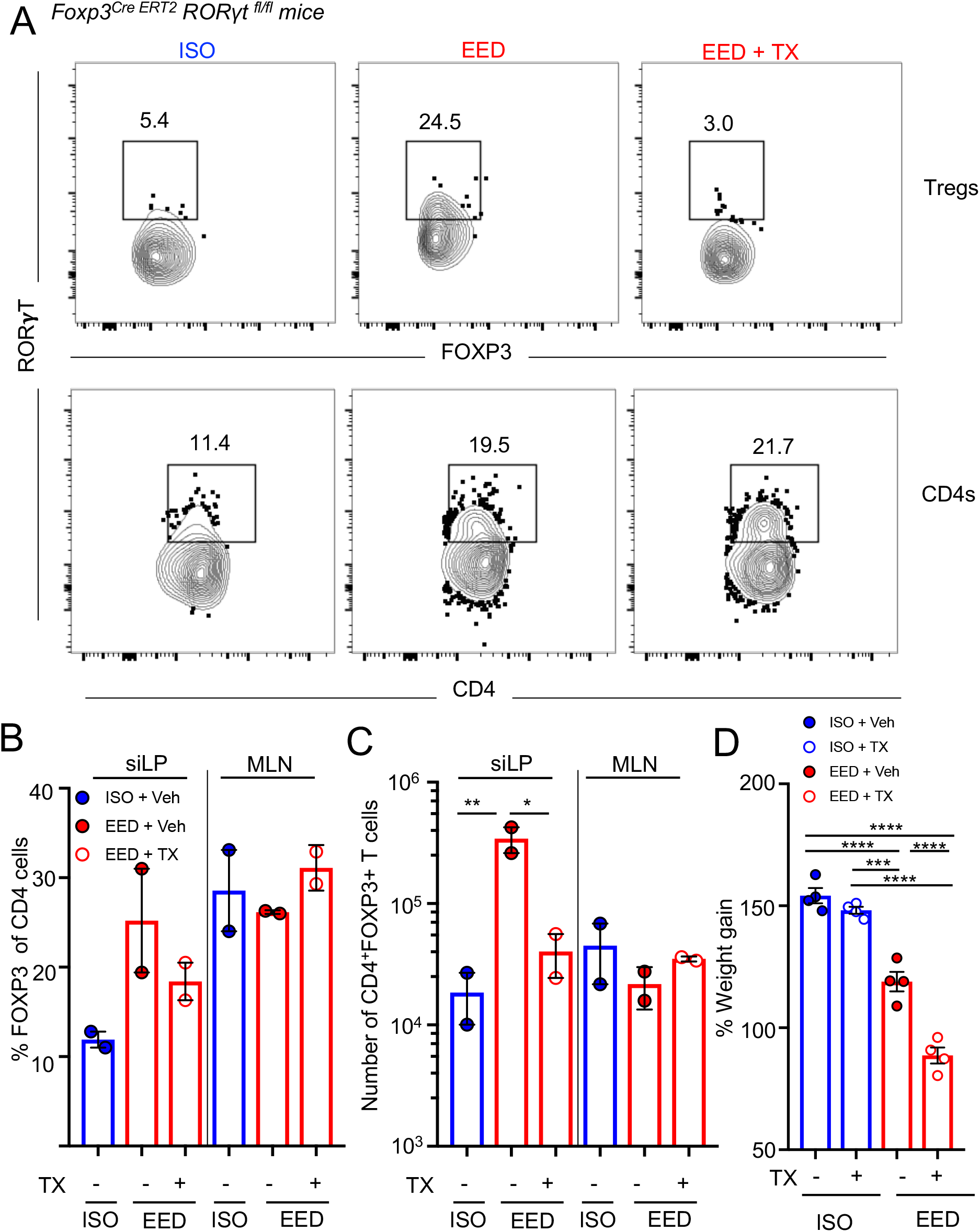
**A**) Representative flow cytometric contour plots showing RORγT expression in FOXP3^+^ CD4^+^ T_regs_ and CD4^+^ T_conv_ cells following treatment of *Foxp3^CreERT2^ Rorc^fl/fl^* mice with tamoxifen, (FOXP3^RORγT-^) or vehicle(corn oil) (FOXP3^RORγT+^). **B**) Percent FOXP3^+^ CD4^+^ T_regs_ after tamoxifen treatment of *Foxp3^Cre ERT2^ RORγt^fl/fl^* mice at day 28 of the EED protocol **C**) Number of FOXP3^+^ CD4^+^ T_regs_ from **B**). **D**) Percent weight gain during EED establishment at day 28 from *Foxp3^Cre ERT2^ RORγt^fl/fl^* mice following treatment with tamoxifen(FOXP3^RORγT-^) or vehicle (FOXP3^RORγT+^) Data points represent a single mouse. Data are represented as mean ± SEM. (p > 0.05; *p < 0.05; **p < 0.01; ***p < 0.001). Data shown is representative of 1-2 experiments.

